# Hydration-dehydration cycles drive compartment dynamics in minimal protocells

**DOI:** 10.64898/2026.03.10.710739

**Authors:** Rafal Zdanowicz, Darshan Chandramowli, Nicola De Franceschi

## Abstract

Compartmentalization is a defining feature of cellular systems, yet how early compartments could undergo repeated cycles of growth, division, and content organization without complex chemistry remains unresolved. Here we study a minimal membrane-based system subjected to periodic hydration– dehydration cycles, mimicking fluctuating physical environments on the early Earth. We show that cyclic environmental conditions alone drive a sequence of reproducible compartment dynamics, including macromolecule encapsulation, membrane growth, division, and the generation of a highly crowded interior. These processes emerge from biophysical transformations of a single-component membrane and do not require any chemical reactions or metabolic activity. Importantly, compartments retain their structural integrity across multiple cycles, enabling repeated encapsulation without loss of individuality. Our results demonstrate that fluctuating physical conditions can be transduced by membrane biophysics into sustained, cell-like cycles, challenging the view that primordial cellular dynamics necessarily required chemically driven growth and division.

## INTRODUCTION

Cellular compartments enable the maintenance of genetic material and the generation of electrochemical gradients, which are key functional requirements of living systems^1,2^. However, the presence of a membranous compartment introduces a series of challenges: cells need to be able to grow, divide, encapsulate content, generate a crowded cytoplasm, and maintain physical integrity. In contemporary cells, these functions are achieved through active processes such as biosynthesis, cytoskeletal force generation, and membrane transport. However, it remains unclear how early cellular compartments could have exhibited repeated cycles of growth, division, and content organization in the absence of such biochemical complexity. This raises a fundamental question: can simple membranes transduce environmental physical fluctuations into sustained, cell-like dynamics without relying on specific chemical reactions?

Most experimental models achieve protocells membrane dynamics by introducing chemically driven processes. Membrane growth is typically induced by the addition of fatty acids micelles^3,4,5,6,7,8^, while division is often coupled to specific reactions or catalytic pathways^9,10^. Although chemically mediated approaches provide valuable insight into the co-evolution of membranes and metabolism, they implicitly assume that early cellular dynamics were controlled primarily by molecular chemistry. This assumption constrains generality: division and growth become contingent on the availability and regulation of particular reactions, rather than emerging as robust physical processes.

A key challenge concerns the coupling between encapsulation and structural integrity. Membrane swelling from a dry state provides an efficient route to encapsulate macromolecules^11,12,13,14,15^, yet it is incompatible with preservation of structural integrity and with maintenance of compartment individuality, leading to loss of compartment individuality across cycles^16^. At the same time, living cells maintain a highly crowded cytoplasm^17,18,19^, with macromolecular concentrations far exceeding those achieved in any protocell models. How repeated encapsulation, crowding, growth, and division could occur within a single system, while preserving compartment identity across cycles, poses an unresolved physical problem.

An alternative view is that early cellular behavior may have been governed by fluctuating environmental conditions. On the prebiotic Earth, compartments would have been exposed to repeated hydration and dehydration events. Such cycles constitute a form of periodic external forcing that acts directly on membranes: whether these fluctuations can be converted into reproducible compartment dynamics, rather than irreversible disruption^13^, remains largely unexplored.

Here we address this problem by studying a minimal membrane system subjected to controlled hydration–dehydration (H/D) cycles. In contrast to classical wet–dry cycles^11-15,20-28^, H/D cycling is defined by the absence of a fully dry state (Fig. S1A), a key distinction with important consequences for membrane integrity and dynamics. We show that H/D cycles drive a reproducible sequence of compartment transformations, including encapsulation, membrane growth, division, and the emergence of a densely crowded interior. In addition, we show that macromolecular crowding modulates division dynamics, revealing a previously unexplored physical role of crowding in minimal compartments. These behaviors arise from biophysical changes in a one-component membrane and do not require any chemical reactions or metabolic energy input. Our results demonstrate that fluctuating physical conditions can be converted by membrane biophysics into sustained, cell-like cycles, revealing a purely physical route to early compartment dynamics.

## RESULTS

### Dehydration results in protocell growth and macromolecule encapsulation

To determine how dehydration affects membrane organization, we subjected a homogeneous suspension of oleic acid (OA) to controlled dehydration. As solvent removal increased the OA concentration, micron-scale vesicular compartments were generated, hereafter referred to as protocells (Fig. S1B). Dehydration therefore acts as a physical driving force for membrane growth, continuously increasing the available membrane area.

Under these conditions, protocells predominantly exhibited elongated, tubular morphologies, increasing their aspect ratio through longitudinal membrane extension (Fig. 1A, Fig. S1C, Movie 1A). Such tubulation reflects a high surface-to-volume ratio, previously observed in fatty acid systems following abrupt increases in OA concentration via addition of micelles^5^. Strikingly, during H/D cycles, tubular protocells spontaneously relaxed into quasi-spherical compartments without any externally applied mechanical perturbation (Fig. 1A, Movie 1B). This abrupt shape transition corresponds to a rapid reduction in surface-to-volume ratio and indicates a non-equilibrium membrane instability induced by dehydration-driven membrane growth. Newly formed spherical protocells appeared more frequently in close proximity to pre-existing protocells, which may function as nucleation points for membrane expansion^3^: as the dehydration progressed, this resulted in the formation of “islands” of protocells (Movie 1C).

**Figure 1:**
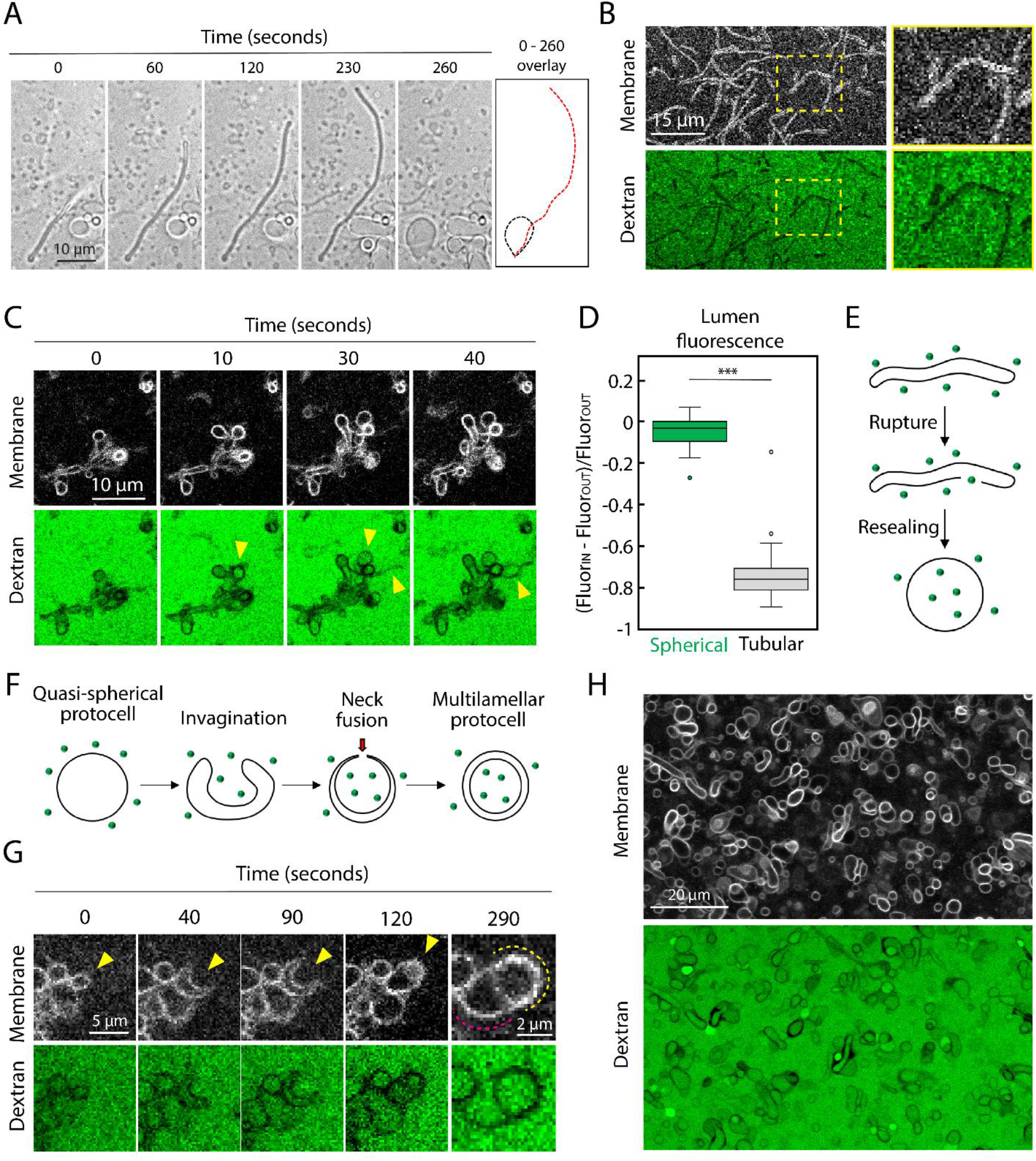
Dehydration drives protocell growth and macromolecule encapsulation. (A): Representative protocell showing tubular growth and collapse into a quasi-spherical morphology during dehydration. (B): Tubular protocells do not display signal from fluorescent dextran in their lumen. (C): Time-lapse frames showing immediate dextran filling coincident with protocell appearance (yellow arrowheads). (D): normalized fluorescence of 150 kDa dextran-FITC in the lumen of tubular protocells (n = 50) and immediately following collapse into a quasi-spherical morphology (n = 77). Bulk fluorescence was defined as zero. Statistical significance was determined by Mann-Whitney U test. (*** p < 0.001). Black horizontal lines indicate median values; boxes represent the interquartile range (25th–75th percentiles), and whiskers extend to 1.5× the interquartile range. (E): Illustration of membrane shape transformation from a tubular to a spherical morphology. Soluble macromolecules are shown as green dots; the membrane is indicated in black. (F): Illustration of membrane shape transformation from a spherical morphology into stomatocyte. (G): Frames from a time-lapse sequence showing a protocell transitioning into a stomatocyte morphology. The magnified view at 290 seconds highlights the increased fluorescence of the double membrane (yellow dotted line) relative to the single outer membrane (red dotted line), confirming stomatocyte formation. (H): Representative field of view during dehydration, showing that the majority of protocells encapsulate dextran.

To test whether this shape transition was accompanied by transient loss of membrane integrity, we performed dehydration in the presence of a hydrophilic macromolecular tracer (150-kDa dextran–FITC). Tubular protocells initially excluded dextran from their lumen (Fig. 1B). However, during the tubular-to-spherical transition, dextran rapidly entered the protocell interior, reaching concentrations comparable to the surrounding bulk solution (Fig. 1C–D, Movie 1C). The coincidence of abrupt shape relaxation and macromolecule entry is consistent with a mechanism involving transient membrane rupture followed by rapid resealing (Fig. 1E). This identifies a previously unrecognized biophysical pathway by which dehydration-driven membrane remodeling enables macromolecule encapsulation without transition through a fully dry state.

In addition to rupture-mediated encapsulation, dehydration activated a second, distinct encapsulation pathway. Increasing osmotic pressure^29^ during dehydration induced inward membrane bending in quasi-spherical protocells, leading to the formation of stomatocyte morphologies (Fig. 1F–G, Movie 1D). Fusion of the stomatocyte neck generated multilamellar compartments (Fig. 1F, Fig. S1D). Accordingly, the average lamellarity of the sample increased during dehydration (Fig. S1E). Stomatocyte formation results in macromolecule encapsulation within internal membrane layers; upon subsequent hydration and shedding of outer lamellae, encapsulated macromolecules become localized within the protocell lumen (Fig. 1F, Fig. S3).

Together, these observations demonstrate that dehydration enables macromolecule encapsulation through at least two physically distinct mechanisms: (i) transient membrane rupture and resealing and (ii) engulfment via dehydration-induced stomatocyte formation. Through the combination of these two mechanisms, the majority of protocells generated during dehydration displayed a lumen containing dextran (Fig. 1H). Importantly, both encapsulation pathways occur concomitantly with membrane growth and do not require protocells to pass through a dry phase, establishing dehydration as a direct physical driver of growth–encapsulation coupling in minimal membrane systems.

### Protocells spontaneously concentrate macromolecules against a concentration gradient

During dehydration, we observed an unexpected phenomenon: a substantial fraction of protocells accumulated macromolecules in their lumen at concentrations exceeding those in the surrounding bulk solution (Fig. 1H, 2A, 2B, 2E, S2A, S2B). These highly fluorescent protocells constituted 20.6 ± 7.9 % of the population (Fig. 2A). The same phenomenon was observed using 10 kDa dextran–Cascade Blue (Fig. S2C) and a fluorescently labelled 23-mer ssDNA (Fig. S2D), demonstrating that macromolecule enrichment is governed by a generic physical mechanism, independent of molecular size or chemical identity. The emergence of this enriched subpopulation was highly reproducible and could be followed in real time during dehydration (Fig. 2B, Movie 2A).

**Figure 2:**
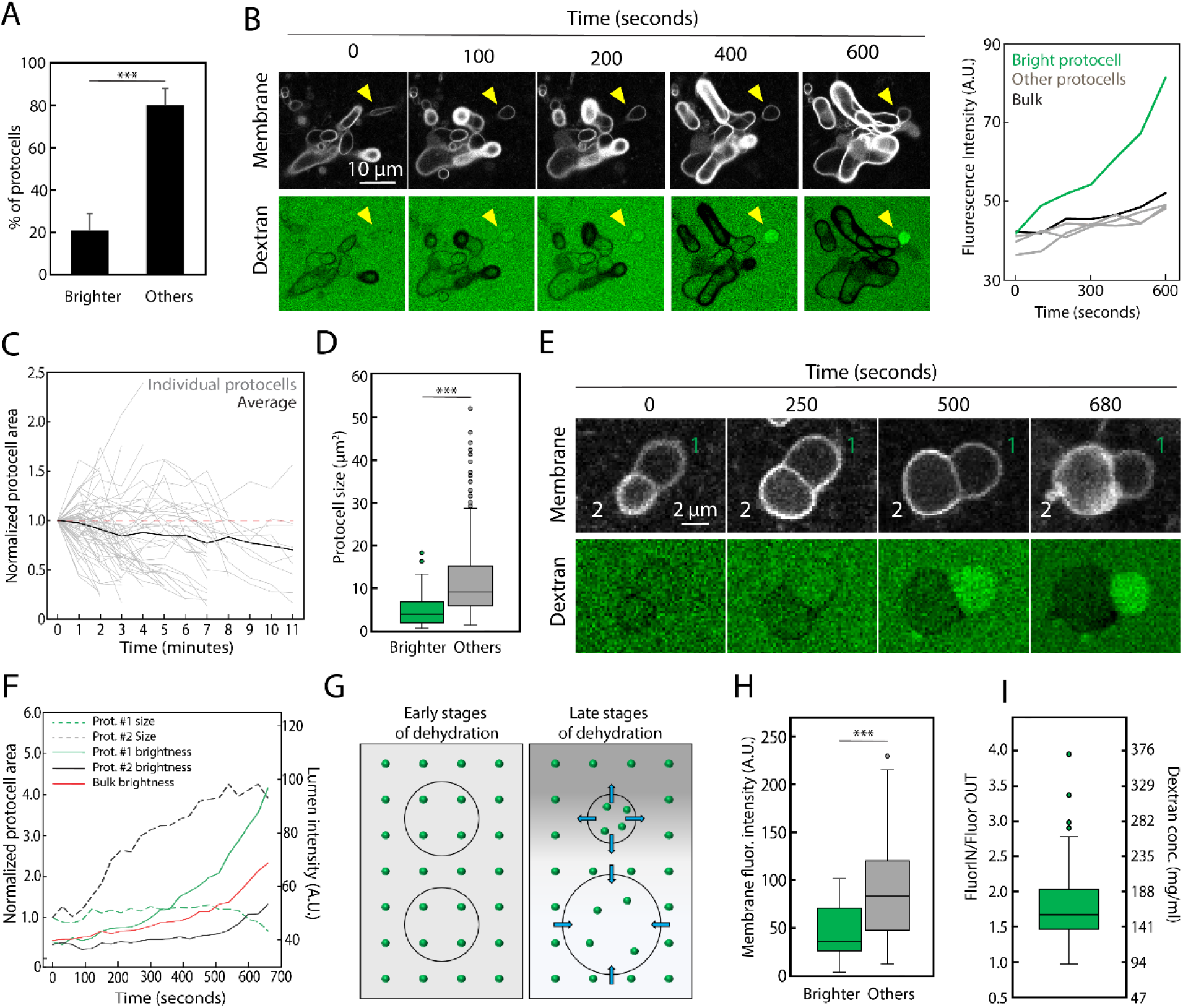
Protocells concentrate macromolecules in their lumen against gradient. (A): Percentage of protocells exhibiting fluorescence intensity above bulk levels in the presence of 150 kDa dextran-FITC. N = 532 and 2253 protocells from 4 independent experiments. Statistical significance was assessed by a 2-tailed T-test. The bars indicate standard deviation. (B): Frames from a time-lapse sequence showing progressive concentration of 150 kDa dextran-FITC within the lumen of a protocell. The overall dextran brightness was adjusted to compensate for the progressive concentration due to dehydration. Quantification of the dextran fluorescence intensity is shown on the right. (C): Quantification of changes in protocells area upon dehydration. N = 62 protocells from 5 independent experiments. (D): Quantification of protocell area upon dehydration. N = 700 protocells from 4 independent experiments. Statistical significance was assessed by a Mann-Whitney U test. (E): Frames from a time-lapse showing concomitant change of size and concentration of encapsulated 150 kDa dextran-FITC. (F): Simultaneous tracking of lumen brightness and volume of the protocells shown in (E). (G): Proposed mechanism for macromolecule concentration inside protocells. Spatially inhomogeneous osmotic pressure, arising from local fluctuations in low–molecular-weight solute concentration (primarily Tris–HCl; grey shaded region), drives water efflux across the membrane (blue arrows), leading to volume reduction and concentration of encapsulated macromolecules (dextran, green dots). (H): Quantification of membrane fluorescence intensity of protocells upon dehydration. N = 35 and 60 protocells from 6 independent experiments. Statistical significance was assessed by a Mann-Whitney U test. (I): Quantification of dextran enrichment inside protocells at the apex of dehydration normalized for the dextran concentration in the bulk, corresponding to 94 mg ml^-1^. n = 166 protocells from 2 independent experiments. In all box-and-whisker plots, black horizontal lines indicate median values; boxes represent the interquartile range (25th–75th percentiles), and whiskers extend to 1.5× the interquartile range. In all plots, ** = p < 0.01 and *** = p < 0.001.

The accumulation of macromolecules inside protocells implies the existence of a mechanism capable of driving concentration against an external gradient. Dehydration progressively increases osmotic pressure in the bulk solution, leading to water efflux from protocell lumens and a reduction in protocell volume (Fig. 2C, Movie 2B). We hypothesized that macromolecule enrichment arises when protocell shrinkage locally outpaces the reduction in bulk solution volume. Consistent with this interpretation, enriched protocells were on average smaller than the rest of the population (4.7 ± 3.2 µm^2^ vs. 12.0 ± 10.3 µm^2^) (Fig. 2D), and simultaneous tracking of protocell size and lumen fluorescence revealed an inverse correlation between volume and macromolecule concentration (Fig. 2E-F). We propose that spatial heterogeneity in osmotic pressure within the dehydrating sample drives differential shrinkage across protocells, resulting in selective macromolecule enrichment in a subset of protocells. Such heterogeneity is plausible given the highly crowded, viscous and dynamically evolving environment experienced by individual protocells during dehydration. In support of this interpretation, we observed that while most protocells underwent shrinkage, others continued to grow under the same global dehydration conditions (Fig. 2C). Macromolecular crowders such as dextran contribute minimally to osmotic pressure, whereas small, abundant solutes dominate osmotic effects. In our system, this role is primarily played by Tris–HCl, present at 200 mM at the beginning of dehydration. This establishes a simple physical mechanism for achieving a high level of crowding: local fluctuations in the concentration of low-molecular-mass solutes generate osmotic pressure differences that concentrate high-molecular-mass components inside protocells (Fig. 2G). In line with this mechanism, enriched protocells exhibited a lower lamellarity compared to non-enriched protocells (45.6 ± 27.8 A.U. vs. 89.7 ± 48.2 A.U) (Fig. 2H). A lower lamellarity renders a protocell more prone to react to variation in osmotic pressure, whereas higher lamellarity insulates it more effectively.

As dehydration approached its culmination, the bulk dextran concentration was estimated to reach 94 mg ml^−1^ (Supplementary Note 1), a regime in which diffusion was strongly suppressed (Fig. S2E-F). Under these conditions, enriched protocells displayed an average 1.76-fold increase in lumen dextran concentration (Fig. 2I), corresponding to 165 mg ml^−1^, comparable to macromolecular crowding levels in living cells^18,19^. The most concentrated protocell reached an estimated dextran concentration of 366 mg ml^−1^. Upon further dehydration, the sample transitioned to a fully dried state, identifiable by a crystal-like morphology (Fig. S2G). In all experiments involving repeated H/D cycling, samples were deliberately prevented from reaching this fully dried state. Together, these results demonstrate that dehydration alone can drive spontaneous enrichment of macromolecules inside protocells against a gradient to physiologically relevant concentrations, without requiring active transport.

### Protocells retain and amplify macromolecular content across H/D cycles

Upon hydration, the protocells clusters formed during dehydration progressively disassembled into individual protocells (Fig. S3A). Cluster fragmentation was likely driven by the combined effects of hypo-osmotic shock, dilution of OA and microscale fluid motion generated during water addition. Although hydration was performed without intentional mixing, turbulence at the microscale is intrinsic to fluid addition and therefore constitutes an integral component of H/D cycles. During hydration, lamellarity decreased through multiple pathways, including membrane rupture (Fig. S3B, Movie 3A) and protocell extrusion events (Fig. S3C, Movie 3B).

Despite these disruptive processes, a subset of protocells retained macromolecules at concentrations exceeding those of the surrounding bulk following hydration. Retention was observed both for samples cycled directly on the microscope stage (Fig. S3D) and for bulk samples cycled in vials (Fig. 3A). Encapsulated dextran persisted for at least 30 h after hydration (Fig. S3E), and similar retention was observed for ssDNA (Fig. S3F) and 10 kDa dextran–Cascade Blue (Fig. S3G). By contrast, control samples subjected to the identical procedure but prevented from dehydrating showed no enrichment relative to the bulk (Fig. 3B), confirming that dehydration is the key physical process enabling enhanced macromolecule encapsulation.

**Figure 3:**
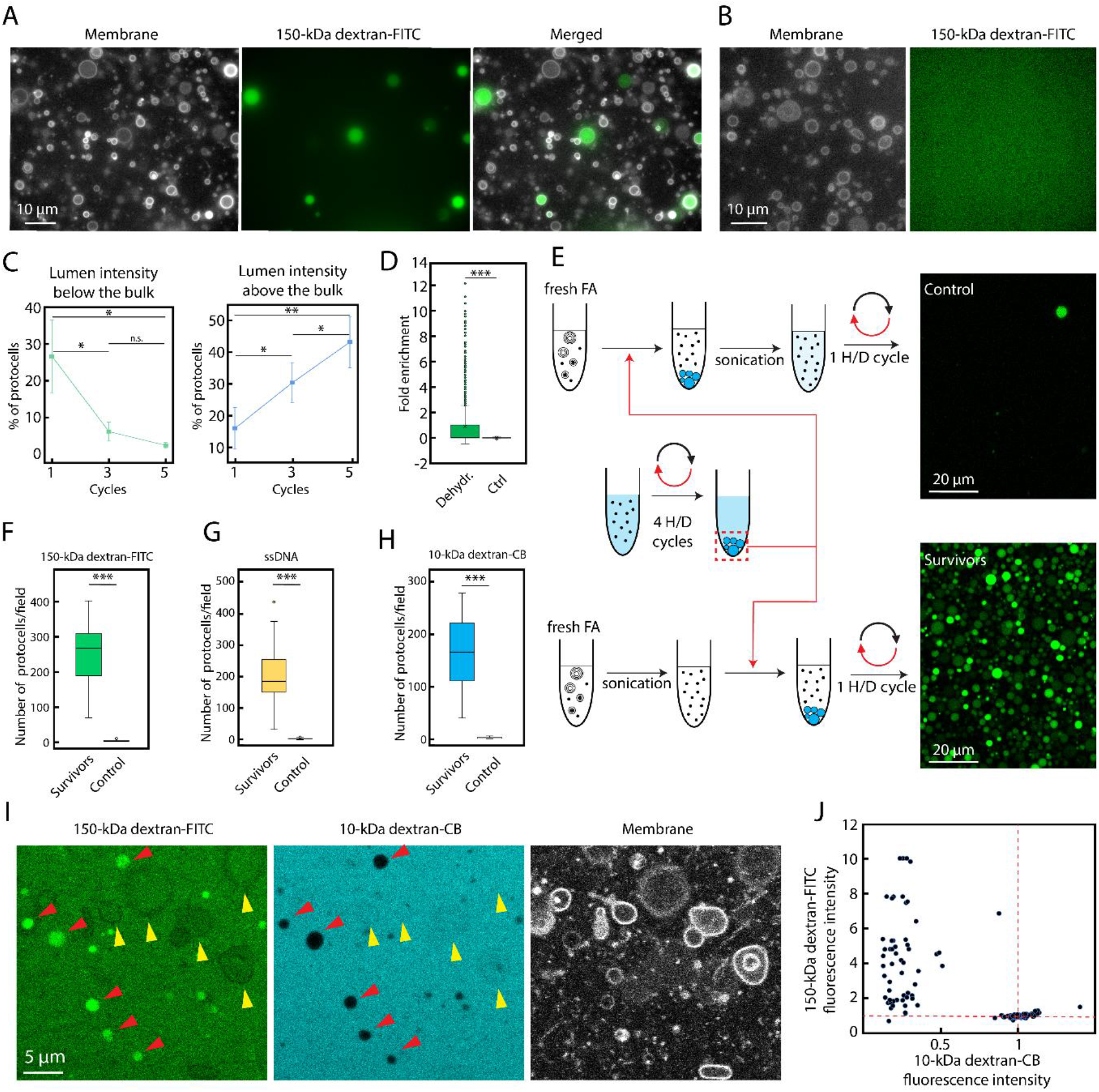
Repeated H/D cycles lead to cumulative macromolecule encapsulation in protocells. (A): Image of protocells obtained after H/D cycling with dehydration, retaining 150-kDa dextran-FITC at a concentration higher than the bulk following hydration. (B) Image of protocells obtained after H/D cycling in the absence of dehydration, showing no dextran retention after hydration. (C): quantification of the percentage of protocells with 150-kDa dextran-FITC intensity below (left) or above (right) the bulk upon repeated H/D cycling. For the left plot, n = 518; for the right plot, n = 1421 from 7 independent experiments. (D): Fold enrichment of dextran within individual protocells relative to the bulk (defined as zero on the y axis) after 5 H/D cycling. n = 3048 from 7 independent experiments. (E): Schematic of the sample preparation workflow used to subject pre-formed protocells to H/D cycling. Representative images for each condition are shown. (F): Quantification of the number of protocells per field of view that survived one H/D cycle while encapsulating 150-kDa dextran-FITC. N = 38 fields and 35 fields from 2 independent experiments. (G): Quantification of the number of protocells per field of view that survived one H/D cycle while encapsulating fluorescently labelled ssDNA. N = 47 fields and 36 fields from 2 independent experiments. (H): Quantification of the number of protocells per field of view that survived one H/D cycle while encapsulating 10-kDa Dextran-Cascade Blue. N = 38 fields and 36 fields from 2 independent experiments. Statistical significance in panels (D), (F), (G), and (H) was assessed by a Mann-Whitney U test. (I): Example of pre-formed protocells encapsulating 150-kDa dextran-FITC (in green) subjected to an additional dehydration phase in the presence of both 150-kDa dextran-FITC and 10-kDa dextran-Cascade Blue (in cyan) in the bulk. Pre-existing protocells do not encapsulate the 10-kDa dextran, while newly formed protocells encapsulate both dextrans. (J): Quantification of dextran fluorescence intensity in the lumen of protocells, as shown in (I). Two populations of protocells, corresponding to pre-existing protocells and newly formed protocells, can be distinguished. The red dotted lines indicate the fluorescence intensity of the dextrans present in the bulk, corresponding to 1 mg/ml. N = 163 protocells from 3 independent experiments. In all box-and-whisker plots, black horizontal lines indicate median values; boxes represent the interquartile range (25th–75th percentiles), and whiskers extend to 1.5× the interquartile range. In all plots, * = p < 0.05; ** = p < 0.01 and *** = p < 0.001.

Motivated by these observations, we subjected protocells to repeated H/D cycles. Strikingly, the fraction of protocells exhibiting elevated macromolecule concentration increased monotonically with cycle number, while the proportion of protocells dimmer than the bulk decreased (Fig. 3C). This progressive shift indicates that H/D cycling does not simply reset the initial conditions, but instead selectively favors protocells capable of retaining macromolecular content.

During dehydration in bulk vials, the sample volume decreased from 500 µl to 176 ± 12 µl, corresponding to a 2.8-fold concentration increase. While most protocells exhibited enrichment within this range, a distinct subpopulation displayed greater than 10-fold macromolecule enrichment (Fig. 3D). These highly enriched protocells cannot be accounted for by bulk concentration alone and are consistent with the spontaneous concentration mechanism presented in Fig. 2.

The increasing abundance of macromolecule-filled protocells across cycles implies that some protocells survive H/D cycling without losing their internal content. To test this directly, we generated dextran-filled protocells by performing four H/D cycles, then transferred them into a fresh, sonicated OA suspension prior to an additional cycle. As a control, an identical sample was sonicated immediately before the additional cycle, disrupting all pre-existing protocells. Because the chemical composition of both samples was identical, any difference in the number of macromolecule-filled protocells after cycling could be attributed unambiguously to the survival of pre-existing compartments (Fig. 3E). Consistent with this expectation, protocells present before the additional cycle were able to persist through dehydration and hydration while retaining their contents (Fig. 3F–H).

In parallel, new protocells are continuously generated during each dehydration cycle. To distinguish between retained and newly encapsulated content, pre-formed protocells containing 150 kDa dextran– FITC were dehydrated in the presence of both 150 kDa dextran–FITC and 10 kDa dextran–Cascade Blue in the bulk solution. Newly formed protocells indiscriminately encapsulated both tracers, whereas pre-existing protocells retained only the original dextran–FITC cargo (Fig. 3I–J). Together, these results demonstrate that H/D cycling simultaneously enables protocell survival, content retention, and the continual generation of new macromolecule-filled compartments, resulting in a cumulative amplification of macromolecule-encapsulating protocells over successive cycles.

### A crowded protocell lumen biases membrane deformation towards division

During dehydration, pre-formed protocells containing either dextran or ssDNA underwent pronounced shape deformations, transitioning from quasi-spherical morphologies to dumbbell-like structures (Fig. 4A). Larger protocells frequently displayed multiple lobes of heterogeneous size (Fig. 4B, Movie 4A), reminiscent of morphologies observed in L-form bacteria^30^. Because physical division requires both membrane deformation and scission^31^, we next asked whether these dumbbell structures corresponded to fully separated compartments.

**Figure 4:**
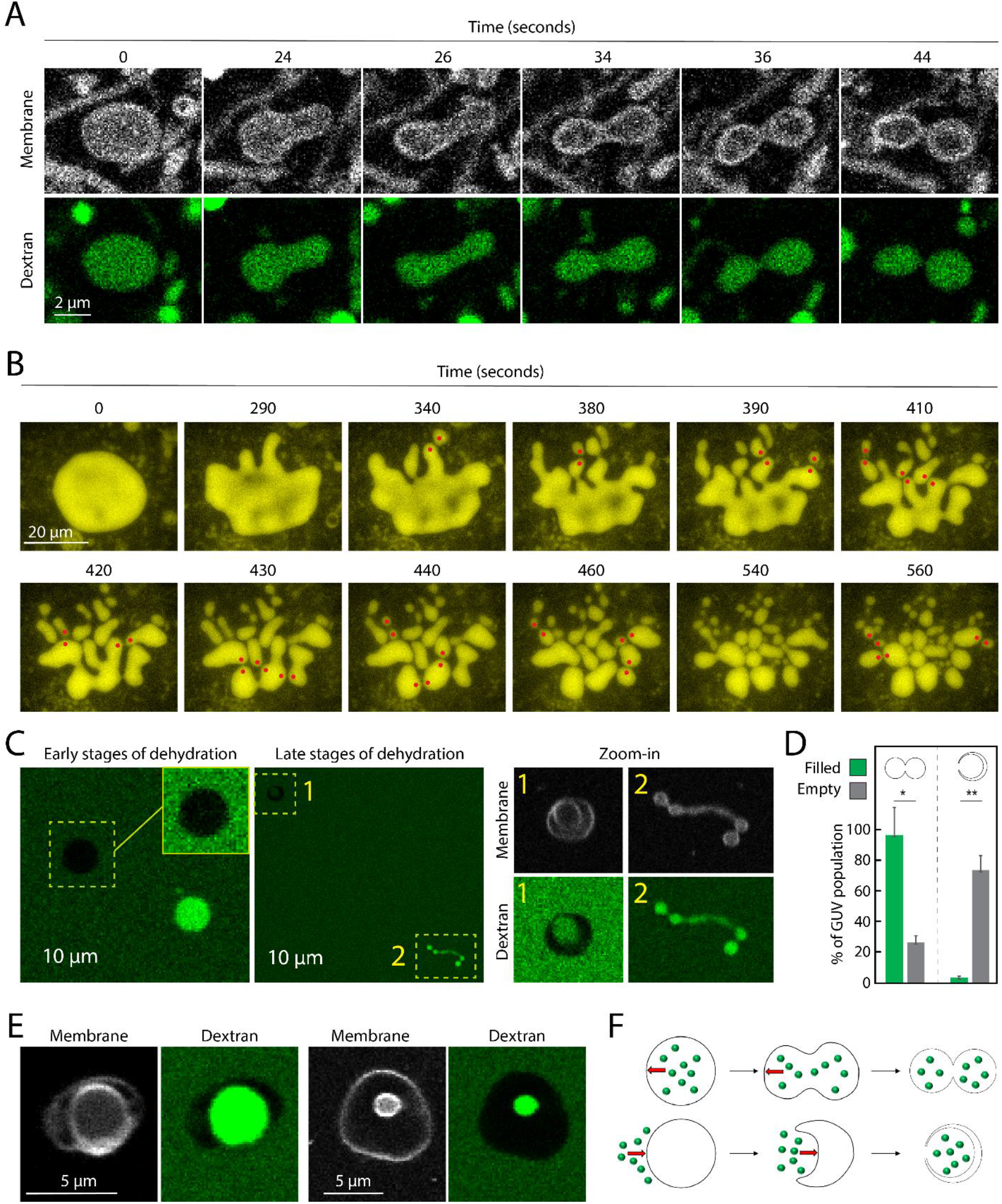
Macromolecular crowding biases protocell deformation and division during H/D cycles. (A): Frames from a time-lapse sequence showing a protocell encapsulating 150-kDa dextran-FITC undergoing a shape transformation into a dumbbell morphology. (B): Frames from a time-lapse sequence showing a large protocell encapsulating fluorescently labelled ssDNA undergoing a shape transformation into multiple dumbbells. The red dots indicate newly formed membrane necks. The ssDNA fluorescence is shown in yellow. (C): Representative examples of spherical GUVs undergoing shape transformations into dumbbell or stomatocyte morphologies upon dehydration. In the magnified views, dextran fluorescence was enhanced post-acquisition to highlight the negative outline of empty GUVs. (D): Quantification of the experiment in (C), showing that GUVs with crowded lumen reshape preferentially into dumbbells. N = 195 filled GUVs and 149 empty GUVs from 7 independent experiments. Statistical significance was assessed by T-test.* = p < 0.05;. ** = p < 0.01. (E): Representative images of empty GUVs engulfing dextran-filled GUVs upon dehydration. (F): Schematic summarizing the role of macromolecular crowding in vesicle deformation. External macromolecules exert physical pressure that drives transitions to stomatocyte, whereas lumen-confined macromolecules induce deformation into dumbbells. Macromolecules are depicted as green dots; the direction of the pressure is indicated by red arrows.

To probe luminal connectivity between lobes, we performed fluorescence recovery after photobleaching (FRAP) experiments^32,33^ using 150-kDa dextran–FITC encapsulated within dumbbell-shaped protocells. No fluorescence recovery was observed following photobleaching of individual lobes (Fig. S5A), indicating that the lumens of adjacent lobes were not in communication. This result is consistent with the presence of either hemi-scission or full scission states. Direct FRAP measurements of membrane connectivity^33^ could not be performed in this system because the membrane dye partitions into the aqueous phase and rapidly re-equilibrates between compartments (Fig. S5B). However, two observations suggest that dehydration alone is generally insufficient to trigger complete scission: elongated membrane necks frequently persisted between lobes (Fig. S5A), and lobes within dumbbell chains remained clustered rather than diffusing independently (Fig. 5B).

**Figure 5:**
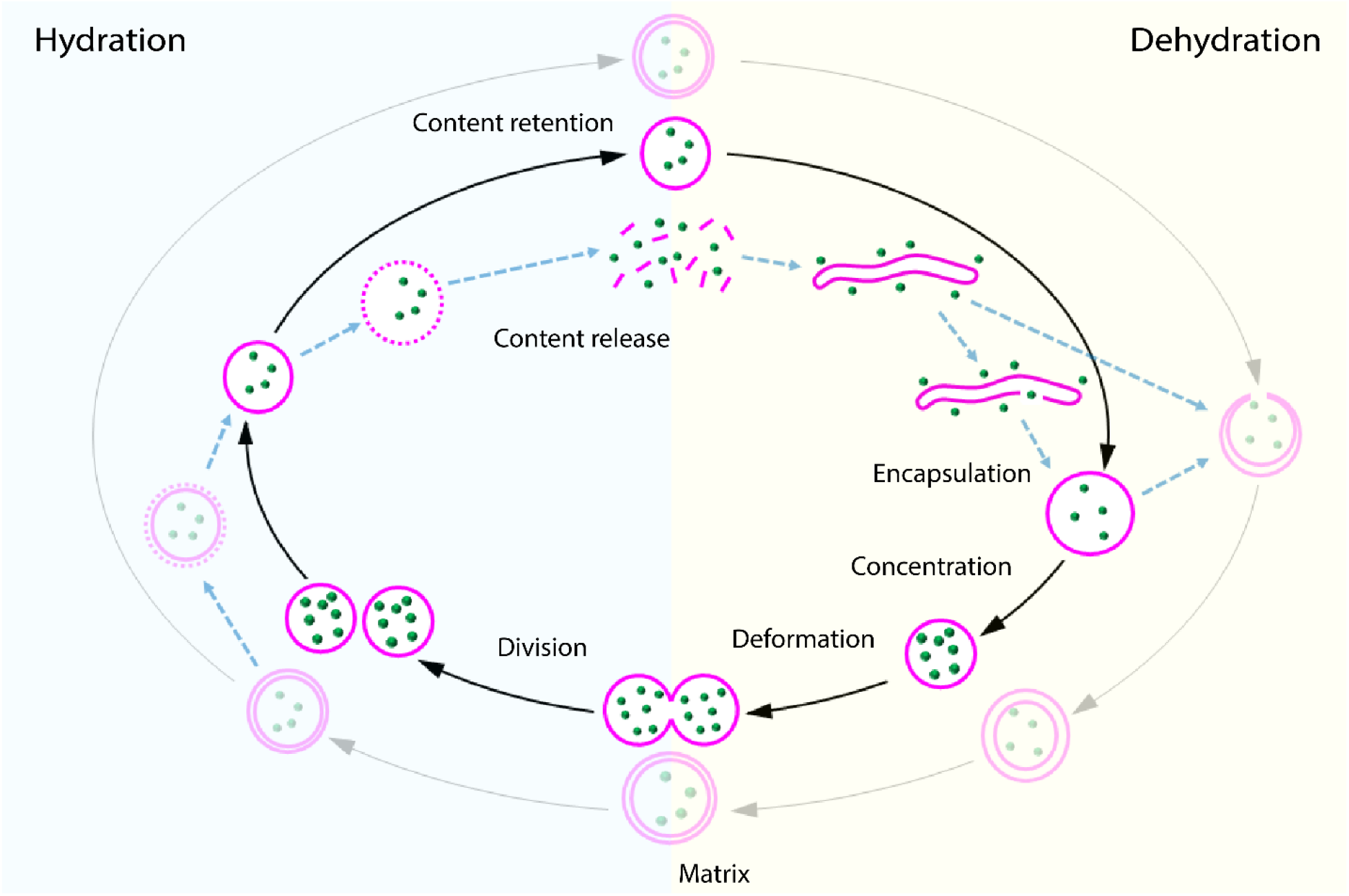
Physical model for hydration–dehydration driven protocell cycles. During dehydration, protocells undergo membrane growth, encapsulation and concentration of macromolecules, and deformation into dumbbell morphologies. Upon hydration, dumbbells divide and macromolecular content is retained despite bulk dilution. Protocell lamellarity increases during dehydration and decreases upon hydration, resulting in the coexistence of protocells with varying degrees of multilamellarity. Complete membrane loss leads to content release. At each cycle, a new generation of protocells re-encapsulates all macromolecules present in the bulk.

These observations point to hydration as the trigger for final scission. Hydration induces hypo-osmotic shock, OA dilution and microscale fluid motion, all of which could trigger scission. Consistent with this interpretation, protocell clusters formed during dehydration dispersed upon hydration, yielding separated individual protocells (Fig. 3A, 3E, 3I, Fig. S3E–G). Together, these results indicate that protocell division emerges spontaneously from H/D cycles, with dehydration driving membrane deformation and hydration providing the mechanical stimulus required for scission.

Notably, the mode of deformation depended strongly on protocell content. Pre-formed protocells containing macromolecules deformed predominantly into dumbbell morphologies (Fig. S5C), whereas newly formed protocells or protocells with low internal content preferentially deformed into stomatocytes (Fig. 1G, 3I). To isolate the physical origin of this asymmetry from the confounding effects of membrane growth and multilamellarity, we turned to a simplified model system consisting of phospholipid giant unilamellar vesicles (GUVs).

GUVs containing 10 mg ml^−1^ dextran in their lumen were mixed with empty GUVs in a bulk solution containing 2 mg ml^−1^ dextran, with osmolarity carefully equalized between populations. At the onset of dehydration, both dextran-filled and empty GUVs appeared spherical (Fig. 4C). As dehydration progressed, empty GUVs deformed predominantly into stomatocytes, whereas dextran-filled GUVs deformed almost exclusively into dumbbell shapes (Fig. 4C–D). This bias was particularly evident in occasional pairs of adhered GUVs: in such pairs, the empty GUV invariably attempted to engulf the dextran-filled GUV, regardless of their relative sizes (Fig. 4E).

Because no detectable binding of dextran to the membrane was observed (Fig. 4A, 4C, 4E), asymmetric membrane interactions^34^ cannot account for the observed directionality. We therefore propose that macromolecular crowding generates a physical pressure that biases membrane deformation away from the crowded side (Fig. 4F). This crowding-induced mechanical asymmetry represents a previously unrecognized physical effect of confined macromolecules on membrane shape.

Within H/D cycles, this mechanism establishes a feedback loop between internal content and division. Newly formed protocells and protocells that failed to enrich macromolecules in previous H/D cycles preferentially undergo stomatocyte deformation and engulf external material, whereas protocells that successfully concentrated crowders preferentially deform into dumbbells and undergo division upon hydration. As a result, H/D cycling couples macromolecular enrichment to division through purely physical mechanisms, without requiring chemical reactions, molecular machinery or active force generation.

## Discussion

Our results show that hydration–dehydration (H/D) cycles can drive a complete sequence of cell-like behaviors—compartment formation, growth, encapsulation, macromolecular crowding, and division— through purely physical mechanisms. In this system, fluctuating environmental conditions act as a source of non-equilibrium conditions, while membrane responses to osmotic effects and crowding determine compartment dynamics. No chemical reactions or active force-generating machinery are required. Instead, repeated cycling rectifies environmental fluctuations into reproducible compartment transformations, establishing a physically grounded route to sustained protocell dynamics.

Hydration and dehydration play complementary roles within this cycle. Dehydration drives membrane growth, volume reduction, encapsulation, spontaneous macromolecular enrichment, and large-amplitude shape deformations, whereas hydration provides mechanical perturbations that complete scission and reset bulk conditions (Fig. 5). This separation of roles explains how deformation, scission, and content retention can coexist in a minimal system without molecular regulation.

A central insight emerging from this work is the active mechanical role of macromolecular crowding, which generates an internal physical pressure that reshapes the membrane’s mechanical response, biasing deformation to toward division-like, dumbbell morphologies. In this way, internal content becomes mechanically coupled to division propensity. This coupling arises generically from confinement and osmotic effects and does not depend on macromolecular identity, identifying crowding as a fundamental physical regulator of compartment behavior.

An important and previously underappreciated aspect of these processes is the role of membrane lamellarity. With rare exceptions^5^, multilamellar architectures have been largely neglected in origin-of-life models^35,36^, which often assume unilamellar protocells^37^. However, vesicles formed under prebiotically plausible conditions (i.e. by swelling from a raw surface) are overwhelmingly multilamellar. In the H/D system studied here, protocells continuously transition between different degrees of lamellarity, with lamellarity increasing during dehydration and decreasing during hydration. Lamellarity thus emerges as a dynamic, environmentally controlled property rather than a fixed structural feature.

These observations outline a more realistic scenario in which early protocell populations were heterogeneous, spanning a range of lamellar architectures. Reduced lamellarity enhances responsiveness to osmotic fluctuations, promotes macromolecular crowding, and increases the likelihood of division. Division is therefore mechanically coupled to low lamellarity through the establishment of a crowded interior, providing a physical bias against highly multilamellar compartments.

An additional consequence of this mechanism is that protocell dynamics do not require a continuous external supply of amphiphiles. H/D cycles operate primarily by redistributing and remodeling existing membrane material rather than relying on constant influx, a feature particularly relevant in prebiotic environments where amphiphile availability was likely limited^38^. More broadly, this work addresses a longstanding challenge in protocell research: how growth, division, permeability, encapsulation, and internal organization could be integrated within a single, prebiotically plausible cyclic system. Previous landmark studies have elucidated molecular mechanisms underlying protocell growth^39-41^, division^5^, and membrane permeability^42^, often in isolation. Here, we show that H/D cycling unifies these processes, which emerge as interconnected outcomes within a single physical framework rather than as independently engineered functions.

Our results suggest a complementary perspective on early cellular organization. Origin-of-life research has traditionally focused on informational polymers, metabolic networks, and the chemical reactions associated with them. While these components are undoubtedly central to life, our findings highlight the equally indispensable roles of boundaries, gradients, confinement, and crowding. In the present system, membranes do not merely enclose chemistry; they actively convert environmental fluctuations into spatial organization and dynamic behavior.

In summary, we present a simple, prebiotically plausible cyclical system in which H/D cycles drive a physically integrated protocell cycle. Through the interplay of osmotic forcing, membrane mechanics, and macromolecular crowding, this system recapitulates key features of living cells while remaining chemically minimal. Future work will exploit these physical features to increase biochemical complexity, including the spontaneous generation of electrochemical gradients across membranes. More generally, our results demonstrate how environmental fluctuations can be transduced into sustained, cell-like organization through generic physical principles, providing a robust foundation for early cellular life.

## Acknowledgments

We thank Magda Konarska for useful discussion. This research is part of the project No. 2022/45/P/NZ1/01565 co-funded by the National Science Centre and the European Union Framework Programme for Research and Innovation Horizon 2020 under the Marie Skłodowska-Curie grant agreement No. 945339. We acknowledge funding support from the NCN Sonata Bis 2022/46/E/NZ1/00160 and NCN OPUS 2025/57/B/ST4/02483 grants.

## Authors contribution

N.D.F. designed the study and the experimental setup; N.D.F. collected all data; R.Z. developed the automatic pipeline for protocell analysis; N.D.F., D.C. and R.Z. analyzed the data; N.D.F. wrote the manuscript with inputs from all authors; N.D.F. acquired funding and supervised the work.

