## Supplementary Figures 1 to 4 for "Hydration-dehydration cycles drive compartment dynamics in minimal protocells"

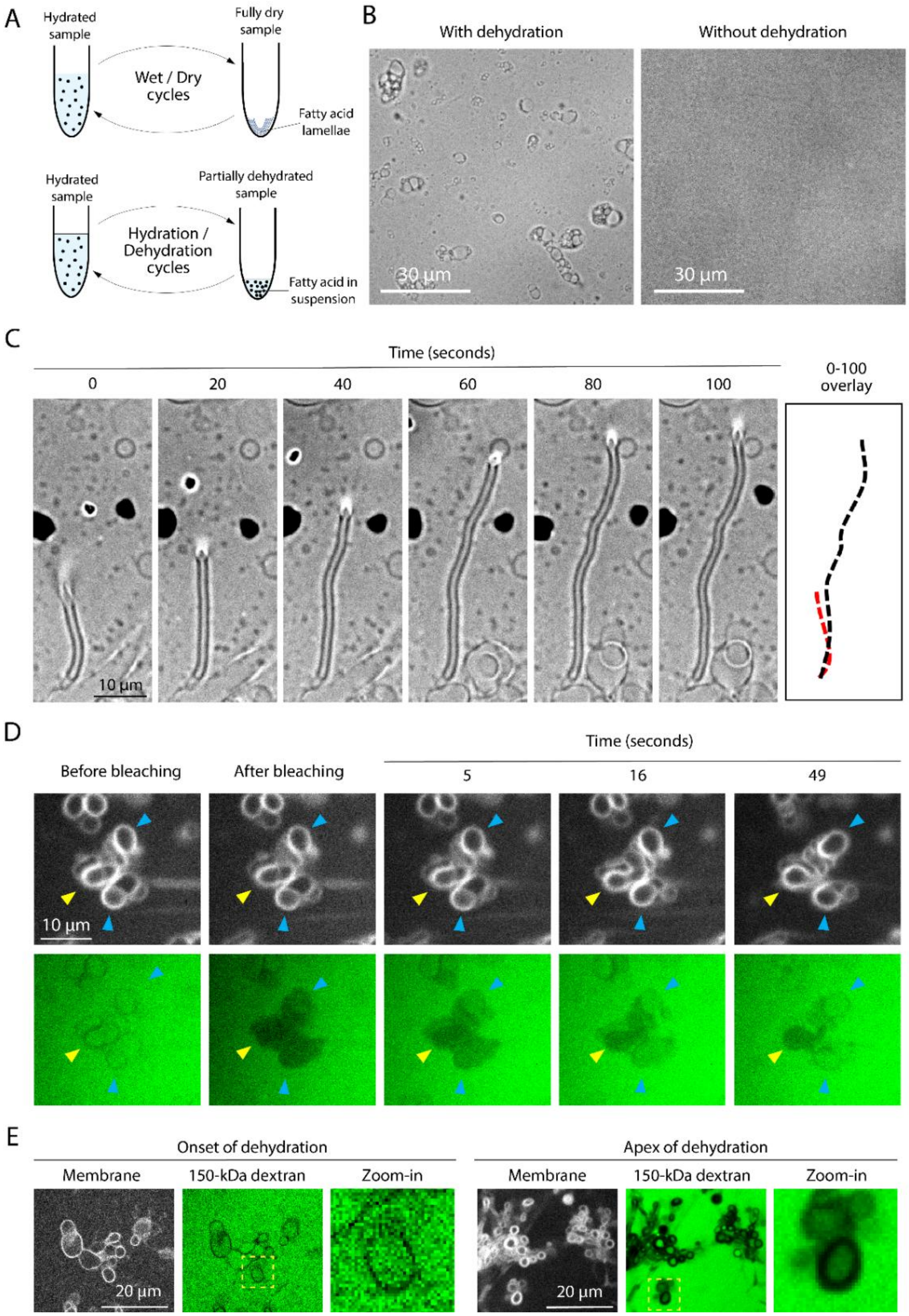


**Supplementary Figure 1: Protocells growth, multilamellarity and content encapsulation.** (A) Schematic comparison highlighting the absence of a fully dry state in hydration–dehydration (H/D) cycles relative to wet–dry (W/D) cycles. (B): Brightfield images of a sonicated oleic acid suspension (4 mM) subjected to dehydration (left) or maintained under sealed conditions to prevent dehydration (right). Dehydration increases the effective oleic acid concentration and induces the formation of microscopic protocells. (C): Frames from time-lapse brightfield microscopy showing tubular growth of a protocell upon dehydration. (D): Fluorescence recovery after photobleaching (FRAP) of a cluster of protocells. Dextran fluorescence within protocells indicated by arrowheads was photobleached, and fluorescence recovery was monitored over time. Protocells marked by blue arrowheads exhibited fluorescence recovery, indicating luminal communication with the external medium (open stomatocytes). In contrast, the protocell marked by the yellow arrowhead showed no recovery, indicating that its lumen was not in communication with the external medium (closed compartment). (E): Representative images of protocells at early and late stages of dehydration, illustrating an increase in lamellarity during dehydration.

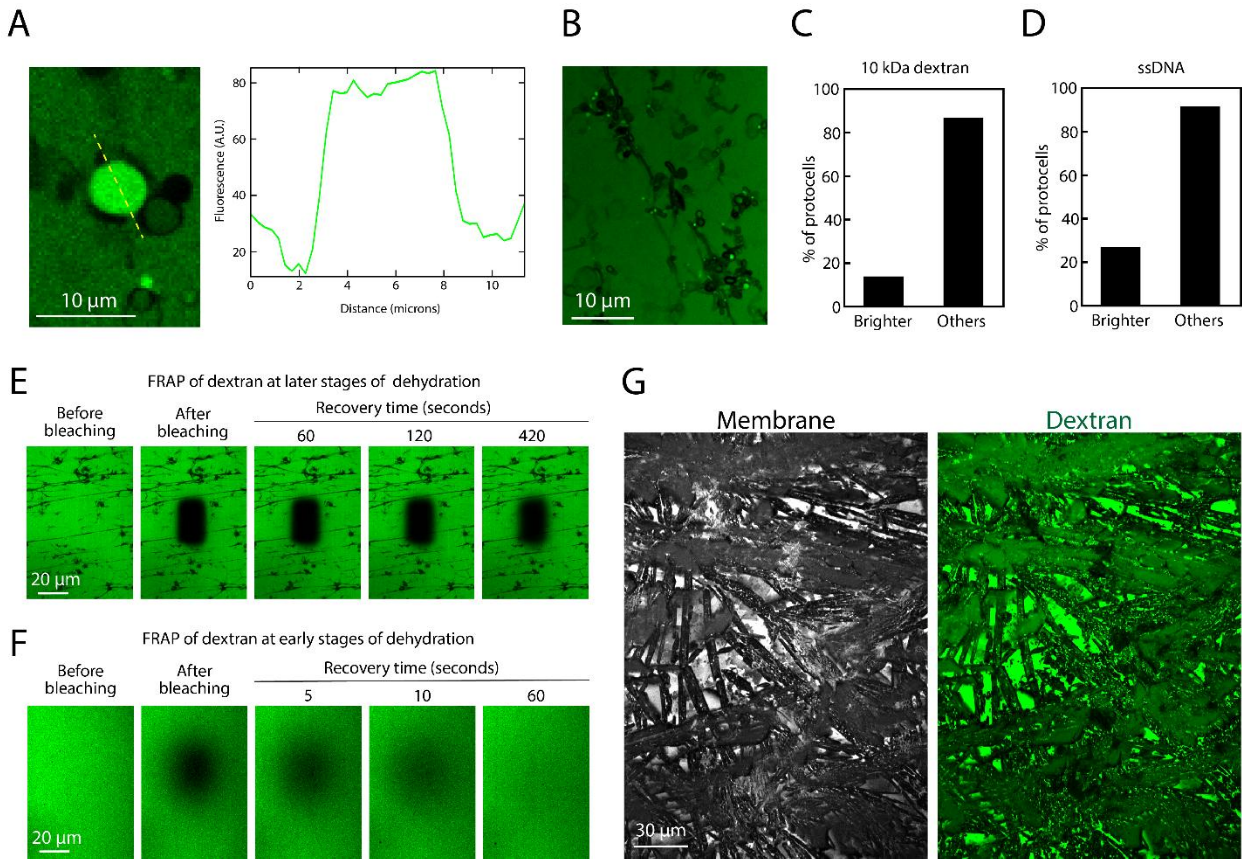


**Supplementary Figure 2: content concentration and environmental conditions during dehydration.** (A, B): Representative protocells with elevated luminal concentrations of 150-kDa dextran relative to the bulk during dehydration. (C): Quantification of the fraction of protocells brighter than the bulk in the presence of 10 kDa dextran–Cascade Blue. N = 980 protocells from 1 experiment. (D): Quantification of the fraction of protocells brighter than the bulk in the presence of Alexa-488-labelled ssDNA. N = 619 protocells from 2 independent experiments. (E) FRAP experiment showing slow diffusion of 150 kDa dextran–FITC at late stages of dehydration. (F) FRAP experiment showing rapid diffusion of 150 kDa dextran–FITC at early stages of dehydration. (G): Representative crystal-like morphology of the protocell sample upon complete drying.

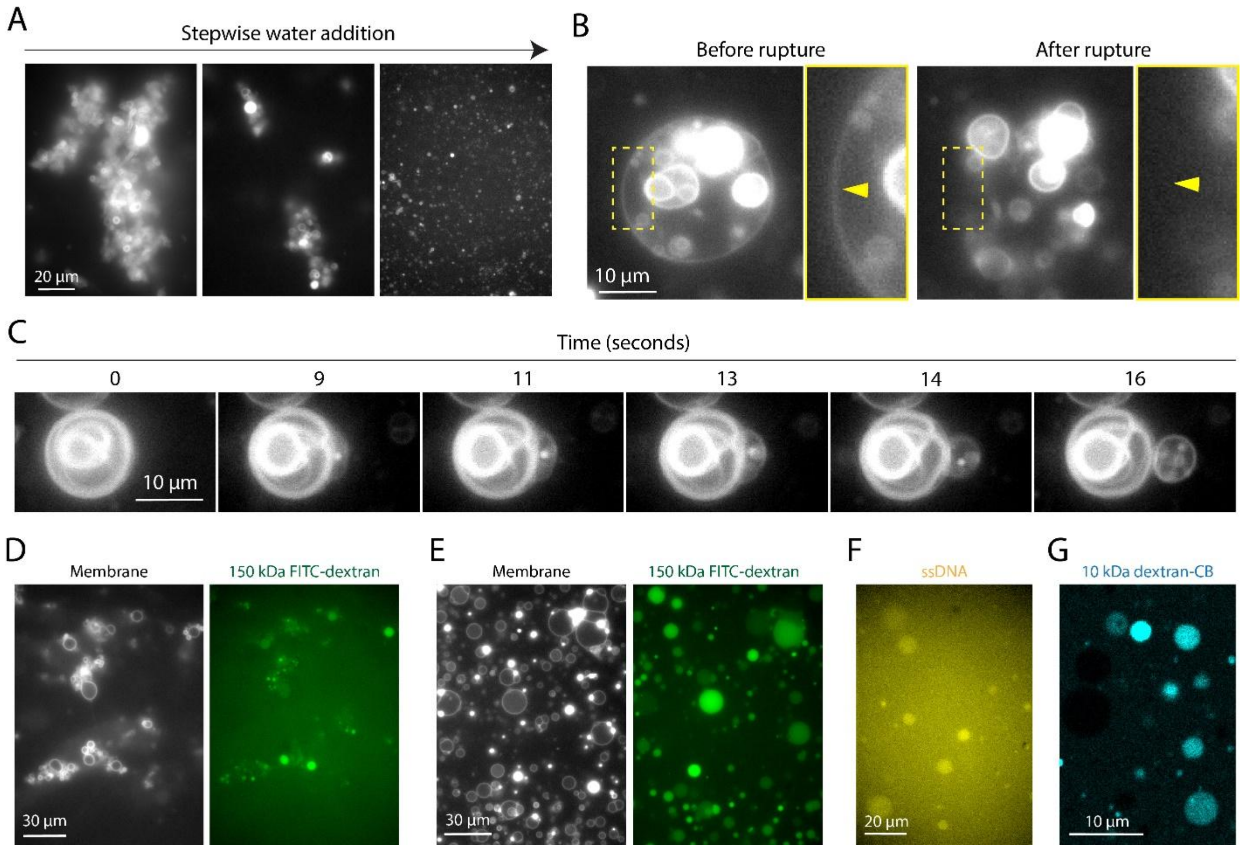


**Supplementary Figure 3: membrane dynamics and macromolecule retention upon hydration.** (A): Representative images illustrating fragmentation of a protocell cluster induced by stepwise addition of water. Membrane fluorescence is shown. (B): Representative example of membrane rupture upon rehydration. The outer membrane (yellow arrowhead in the magnified view) disassembles and eventually disappears. Images show consecutive frames from a time-lapse recording of the same protocell. (C): Protocell extrusion induced by rehydration. Consecutive frames from a time-lapse recording of the same protocell are shown. (D) Representative image of protocells after hydration on the microscope stage, showing retention of dextran at a higher concentration than in the bulk solution. (E) Representative image of protocells after hydration followed by 30 h incubation at 23 °C, showing sustained retention of dextran within the lumen. (F) Representative image of protocells after hydration, retaining fluorescently labelled ssDNA within the lumen. (G) Representative image of protocells after hydration, retaining 10 kDa dextran–Cascade Blue within the lumen.

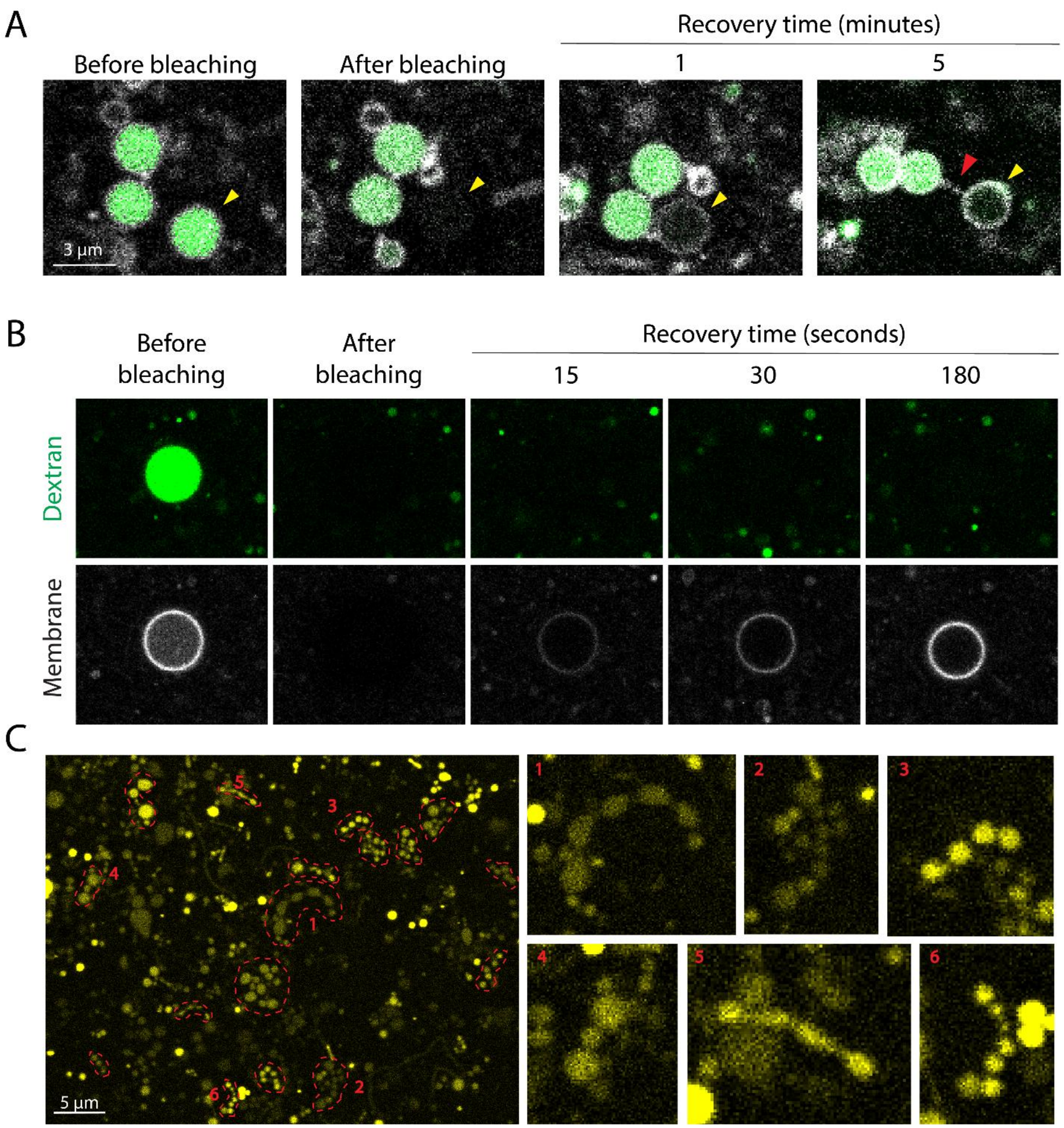


**Supplementary Figure 4: protocell deformation into dumbbells during dehydration.** (A) FRAP experiment in which a lobe of a dumbbell-shaped protocell encapsulating 150 kDa dextran–FITC was photobleached (yellow arrowhead). Fluorescence recovery of dextran was monitored over time, and no recovery was observed. The red arrowhead indicates a membrane nanotube connecting the lobes. (B) FRAP experiment in which both the membrane dye and 150 kDa dextran were photobleached in a spherical protocell. While the membrane dye fluorescence recovered through exchange with the bulk solution, no recovery was observed for dextran. (C) Large field-of-view confocal image of protocells during dehydration, showing widespread deformation into dumbbell-shaped morphologies. Fluorescence from encapsulated, fluorescently labelled ssDNA is shown in yellow.
